# Coiled-coil interactions drive ectopic condensation of overexpressed MAD1 to promote mitotic slippage in cancer cells

**DOI:** 10.1101/2025.07.29.667513

**Authors:** Jason Tones, Jasper Jeffrey, Rachel M. Lackner, David M. Chenoweth, Huaiying Zhang

**Affiliations:** Department of Biology, Carnegie Mellon University, Pittsburgh, PA 15213, USA; Department of Chemistry, University of Pennsylvania, Philadelphia, PA 19014, USA

## Abstract

The mitotic checkpoint protein MAD1 is significantly overexpressed in several cancers, weakening the checkpoint and promoting mitotic slippage. Overexpressed MAD1 forms ectopic foci in mitotic cells, yet the biophysical nature of these foci and their contributions to mitotic slippage remain unclear. Here, we report that MAD1 foci are phase-separated condensates that shorten the mitotic duration by sequestering checkpoint proteins. Our biophysical quantifications reveal that MAD1 ectopic foci in mitotic cells exhibit dynamic condensate properties rather than those of a solid aggregate. Using an inducible phase separation assay in live cells, we show that MAD1 phase separation is driven by interactions between its coiled-coil and disordered domains at the N-terminus. We decouple the contributions of condensation from concentration by inducing the formation of MAD1 ectopic condensates in mitotic cells with low levels of MAD1, demonstrating that the condensation process directly drives mitotic slippage. Mechanistically, the MAD1 ectopic condensate traps the diffusive pool of MAD2, an interaction partner of MAD1, thereby weakening the MAD2 conversion cycle necessary for a robust mitotic checkpoint. Our work illustrates a loss of function caused by ectopic condensates in MAD1-overexpressed cancer cells.

## Introduction

Mitotic arrest deficient 1 (MAD1) is an essential protein in the spindle assembly checkpoint (SAC). The SAC, also known as the mitotic checkpoint, serves as a wait signal to delay chromosome separation until all chromosomes are properly attached to the spindle microtubules in mitotic cells (1, 2). The SAC ensures equal separation of chromosomes into daughter cells, preventing aneuploidy and related diseases such as cancer (3). In a complex with its partner MAD2, MAD1 facilitates the localization of diffuse MAD2 to unattached kinetochores (4). The MAD1-mediated localization of MAD2 at the kinetochore is necessary and sufficient for activating the SAC (5, 6). At the kinetochore, MAD1 promotes a conformational change in MAD2 from its open (MAD2O) form to its closed (MAD2C) form, leading to the formation of the MAD2/CDC20 complex, a crucial step in producing the mitotic checkpoint complex (MCC). The MCC, comprised of MAD2, CDC20, Bub3, and BubR1 in human cells, delays anaphase onset by inhibiting the anaphase-promoting complex (APC/C), which ubiquitinates mitotic proteins such as Cyclin B1 and Securin for degradation (1, 2). Notably, MAD2 is more abundant than MAD1, and the MAD1-free pool of MAD2 is essential for the SAC (7). Besides MAD2, MAD1 also interacts with other mitotic proteins, such as Bub1 and CDC20, and these interactions contribute to robust SAC signaling (8, 9).

Given the significant roles of MAD1 in the spindle assembly checkpoint (SAC), it is not surprising that abnormalities in MAD1 are linked to various cancers (10). Notably, MAD1 overexpression is observed at both mRNA and protein levels in human breast cancers (11–13).

A survey of breast cancer patient tissues indicates that most exhibit high levels of MAD1 protein, and MAD1 overexpression is also associated with a poor prognosis for cancer patients treated with the chemotherapy drug Taxol, which induces mitotic death by targeting spindle microtubules to maintain a persistent SAC (12). Another study using quantitative immunofluorescence microscopy shows that 60% of breast tumors express fivefold more MAD1 protein than normal samples, while 44% express 35-fold more than normal tissues (13).

Mechanistic investigations reveal that MAD1 overexpression decreases MAD2 levels at the kinetochore, weakening the SAC (13). Consequently, cells with overexpressed MAD1 fail to arrest in mitosis when exposed to microtubule poisons. Instead, they undergo mitotic slippage, exiting mitosis prematurely to evade mitotic death and thereby develop resistance to microtubule chemotherapy treatment (13).

The role of overexpressed MAD1 in cancer progression has been linked to its abnormal localization pattern. In interphase cells, endogenous MAD1 is found at the nuclear envelope (14). Along with localizing to the nuclear membrane, overexpressed MAD1 also forms foci in interphase cells. Some ectopic foci contain Nup93, a member of the nuclear pore complex, and resemble a membrane-containing structure known as annulate lamella in the cytoplasm (14). Ectopic MAD1 foci in the nucleus are located at promyelocytic leukemia (PML) bodies (15). This localization displaces MDM2 from PML bodies, enabling MDM2 to ubiquitinate the tumor suppressor p53 for degradation, thus promoting tumor formation (15). In mitotic cells, endogenous MAD1 forms foci on unattached kinetochores (14). In addition to foci on unattached kinetochores, overexpressed MAD1 also forms ectopic foci in the mitotic cytoplasm (13). However, unlike the ectopic MAD1 foci in interphase cells, the biophysical characteristics and disease relevance of these ectopic MAD1 foci in mitotic cells remain unclear.

In this study, we characterize the biophysical properties of ectopic MAD1 foci in the mitotic cytoplasm using quantitative microscopy. Additionally, we investigate the role of MAD1 ectopic foci in mitotic slippage through small molecule-induced MAD1 foci formation. Our findings reveal that ectopic MAD1 foci are not solid aggregates but rather dynamic condensates that can fuse with each other, albeit slowly. Using an inducible phase separation assay in cells, we demonstrate that interactions among the coiled-coil domains and the disordered region at the N-terminus drive MAD1 phase separation. By employing a chemical dimerization system to induce MAD1 phase separation in cells with low MAD1 levels, we show that MAD1 ectopic condensation is sufficient to initiate mitotic slippage in these cells. The toxicity associated with MAD1 ectopic condensation stems from the sequestration of its interacting partner protein MAD2, but not Cyclin B1/CDK1, within the ectopic condensates.

## Results

### MAD1 ectopic foci are dynamic condensates formed in mitotic cytoplasm

We first investigated the biophysical properties of ectopic MAD1 foci in mitotic cells. We transfected exogenous MAD1-mNeonGreen into a HeLa cell line and compared the fluorescence intensity in this line with that of endogenous MAD1 labeled with mNeonGreen through CRISPR knock-in (16). This allowed us to use mNeonGreen fluorescence intensity to quantify the level of overexpressed MAD1 protein at the single-cell level (Fig. 1 A). We added nocodazole to cells released from CDK1 inhibition to arrest them in mitosis and generate unattached kinetochores. Live imaging revealed that at low expression levels, the overexpressed MAD1-mNeonGreen localized to the chromosome in a pattern indicative of unattached kinetochores, similar to the endogenous MAD1-mNeonGreen (Fig. 1A). At high expression levels, in addition to the unattached kinetochore pattern on chromosome DNA, ectopic MAD1 foci that were not localized to chromosomal DNA were observed (yellow arrows, Fig. 1A). The localization patterns are consistent with previous immunofluorescence results from cancer tissues exhibiting various MAD1 expression levels (13).

**Figure 1.**
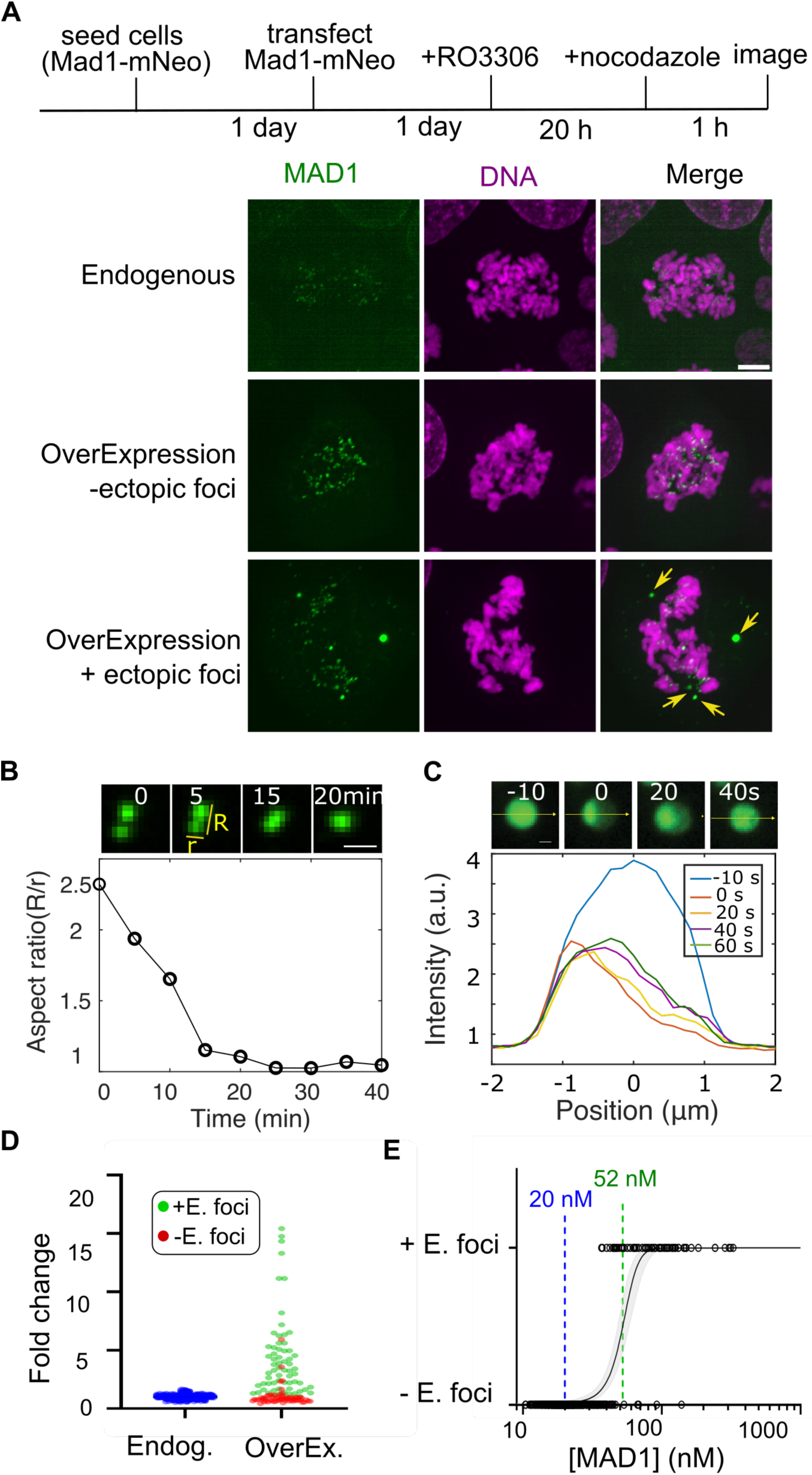
Ectopic MAD1 foci in mitotic cells are dynamic condensates. (A) Experiment schematic and images for nocodazole-arrested mitotic cells with different levels of MAD1 overexpressed. Top panel: HeLa cells with endogenous MAD1 labeled with mNeonGreen. Bottom two panels: HeLa cells with MAD1-mNeonGreen overexpressed via transfection. Yellow arrows indicate ectopic MAD1 foci. (B) Fusion event of ectopic MAD1 foci and the reduction of aspect ratio over time. (C) Images and intensity (yellow line) profile after bleaching half of one ectopic MAD1 focus. (D) Fold change of MAD1 overexpression for cells with/without ectopic MAD1 foci. Each circle represents one cell. (E) Threshold concentration (green line) for the formation of MAD1 ectopic foci. The blue line represents endogenous MAD1 concentration in HeLa cells. Each circle represents one cell. The black line is a logistic regression fit, with gray shaded areas representing the 95% confidence interval. Scale bar in A, 5 μm; Scale bar in B, C, 2μm.

The ectopic foci appear larger and brighter than those on unattached kinetochores. Additionally, unlike the small foci, which remain relatively still, these larger foci exhibit faster mobility in the cytoplasm (Movie 1), likely because they are not attached to the chromosome. The mobile ectopic foci in the mitotic cytoplasm occasionally collide with one another (Movie 1). After a collision, these foci can fuse (Fig. 1B). However, the time required for rounding up to complete fusion is on the scale of minutes, suggesting that these foci have slow dynamics.

Similarly, when half of the MAD1 foci were bleached, fluorescent recovery was observed (Fig. 1C). Again, the time scale of recovery was seen to be on the order of minutes, indicating the slow internal exchange of MAD1 foci. Therefore, we conclude that these ectopic MAD1 foci are not solid aggregates but dynamic condensates formed off kinetochores. Note that because we were unable to observe the fusion of MAD1 foci on kinetochores, we lack sufficient evidence to comment on their physical properties.

Quantifying the mNeonGreen fluorescence intensity and correlating it with ectopic foci formation confirms that MAD1 ectopic foci form at expression levels higher than the endogenous level (Fig. 1D). In addition, only 24% of the cells have MAD1 overexpressed more than five times the endogenous level (Fig. 1D). Considering that 60% of breast tumors express five times more MAD1 protein than normal samples (13), we conclude that our MAD1 expression levels are within the range observed in these tumors.

Based on the reported MAD1 protein concentration of 20 nM in HeLa cells and the 72% labeling efficiency of the CRISPR knock-in (16, 17), we used the average endogenous MAD1 mNeonGreen fluorescence intensity for calibration (Fig. 1E, blue) and converted the fluorescence intensity into MAD1 concentration. We then plotted cells with and without ectopic foci against MAD1 concentration, obtaining the threshold concentration for MAD1 ectopic foci formation at 52.3 ± 2.43 nM, which is 2.6 times the endogenous MAD1 level (Fig. 1E). Together, these data support a threshold behavior typical of phase separation for Mad1 condensate formation.

### Ectopic MAD1 condensates in mitotic cells can form independent of NUP93 and PML

Since MAD1 foci in interphase cells localize to PML bodies in the nucleus and Nup93-labeled annulate lamella in the cytoplasm (14, 15), we next investigated whether ectopic mitotic MAD1 foci co-localized with PML and Nup93 in mitotic cells. Using immunofluorescence with a NUP93 antibody, we observed little co-localization with MAD1 foci in mitotic cells (Fig. S1A), suggesting that mitotic MAD1 ectopic foci are less likely to represent annulate lamella. In a cell line where PML was labeled with the fluorophore Clover through CRISPR, we found that most MAD1 foci formed after MAD1 overexpression did not co-localize with PML foci in mitotic cells (Fig. S1B). This is likely because PML bodies are known to lose most components during mitosis (18). When overexpressing MAD1 in a cell line where PML was deleted via CRISPR, we also observed ectopic MAD1 foci (Fig. S1C), further confirming that MAD1 can form ectopic foci in mitotic cells without PML. Based on these results, we conclude that MAD1 ectopic foci in mitotic cells likely arise from MAD1’s inherent ability to undergo phase separation in the mitotic cytoplasm when overexpressed.

### MAD1 can undergo global phase separation in the presence of local binding sites

The observation that overexpressed MAD1 forms ectopic condensates while also localizing to unattached kinetochores indicates that MAD1 can undergo global phase separation in the presence of local binding sites (Fig. 2A). To directly test this, we adopted the widely used LacO system to mimic MAD1 local binding. In this system, a LacO array was integrated into a U2OS cell line. The fusion of MAD1 to LacI, which binds to LacO, tethers MAD1 to the LacO array to mimic local chromatin binding for MAD1 (Fig. 2A). However, instead of directly fusing MAD1 to LacI, we used a chemical dimerization system to control MAD1 localization to the LacO array, as this method provides the temporal resolution needed to follow the resulting MAD1 phase behavior (Fig. 2B). The dimerizer, Trimethoprim-Fluorobenzamide-Halo ligand (TFH) (19), consists of linked Trimethoprim and Halo ligands that can bind to the eDHFR protein and Halo enzymes, respectively. After fusing MAD1 to eDHFR and Halo to LacI, adding the dimerizer enables MAD1 to dimerize with LacI, thus localizing to the LacO array. Linking MAD1 to LacI, a tetramer (20), increases the interaction valence between MAD1, making it more prone to phase separation at lower concentrations and thus allowing us to follow the condensation process.

**Figure 2.**
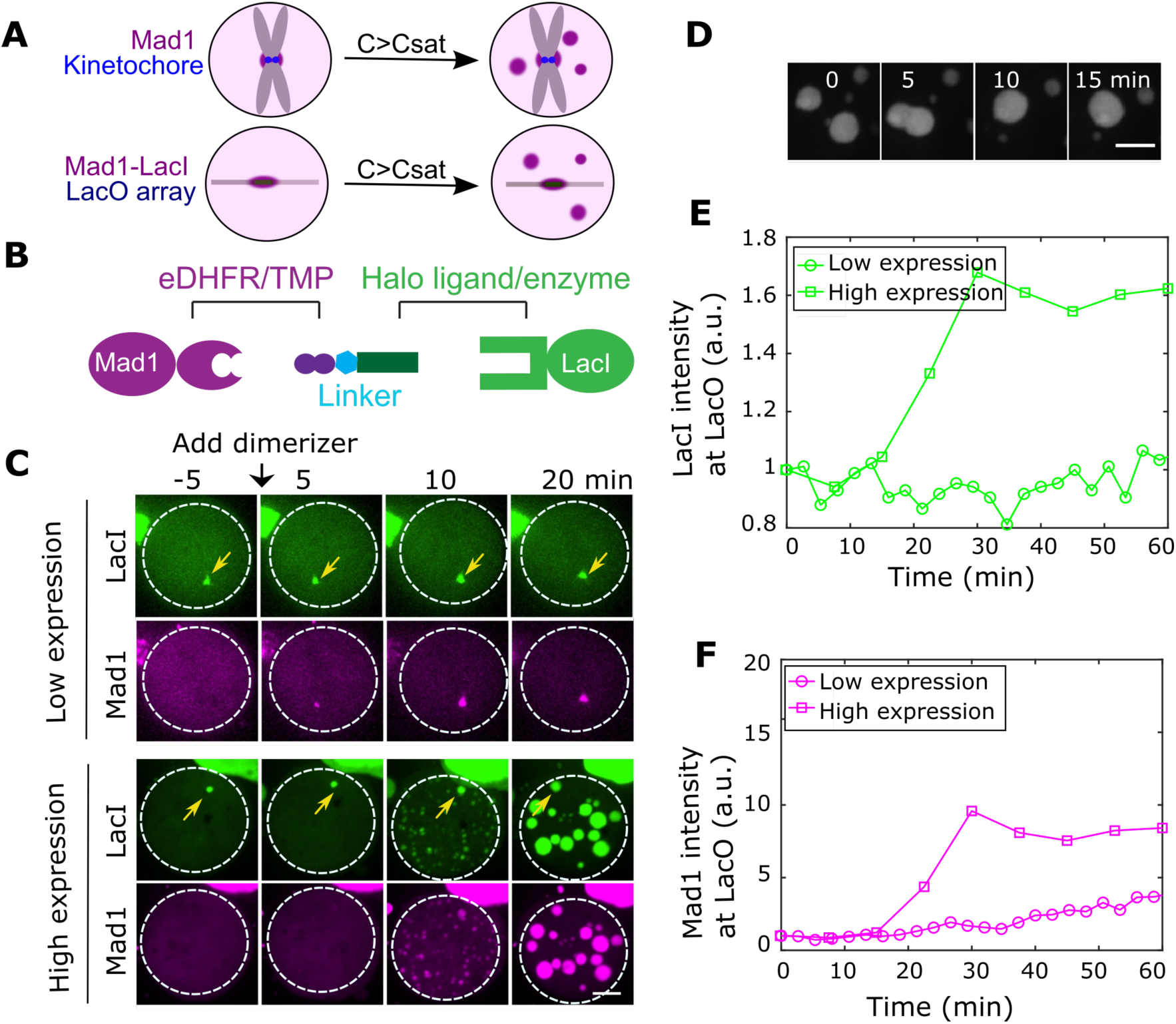
MAD1 can undergo global phase separation in the presence of local binding sites. (A) Schematics using LacO array to mimic MAD1 local binding to kinetochores. (B) Chemical dimerization system and (C) images of recruiting MAD1 to LacO array in mitotic HeLa cells. Cells with MAD1 overexpressed, but without ectopic MAD1 foci were used for this assay. Yellow arrows indicate the LacO array. Dashed lines indicate mitotic cell outlines. (D) Fusion of MAD1-LacI condensates. (E) and (F) quantification of MAD1 and LacI intensity at the LacO array. Scale bar in C, 5 μm; Scale bar in D, 2μm.

We controlled the expression levels of MAD1 to remain below the concentration necessary for spontaneous ectopic condensate formation in mitotic cells (Fig. 2C). Upon the addition of the dimerizer, MAD1 was recruited to the LacO array, as indicated by the increase in MAD1 intensity at the array (yellow arrows, Fig. 2C). However, in cells with relatively high expression levels, global phase separation was triggered, evident from the formation of MAD1 foci in the cytosol (Fig. 2C). The fusion of MAD1 foci in the cytosol was observed, demonstrating dynamic behavior (Fig. 2D). Similar to the spontaneous MAD1 condensates formed through overexpression, the fusion process also took several minutes, suggesting slow dynamics in these induced condensates. Furthermore, the intensity of LacI foci at the LacO array remained constant after dimerization for cells with low MAD1 expression but increased over time for cells with high MAD1 expression (Fig. 2E, F). This increase occurs because high levels of MAD1 also undergo condensation at the LacO array, leading to the localization of MAD1 dimerized-LacI to the LacO array beyond direct LacI-LacO interaction. These findings support that, in addition to local binding, MAD1 can undergo global phase separation in the cytoplasm at sufficiently high concentrations.

### MAD1 phase separation is mainly driven by the coiled-coil domain at its N-terminus

Next, we asked which domains in MAD1 contributed to its phase separation. To achieve this, we used our dimerization system to dimerize MAD1 into a higher valence oligomer, a hexamer HOTag3 (21), allowing it to induce MAD1 phase separation at low levels and be sensitive to changes in the phase behavior of MAD1 truncations (Fig. 3A). First, we verified that the full-length MAD1 can form condensates using this method (Fig. 3B) and that these condensates possess the ability to fuse (Fig. 3C), similar to the spontaneous MAD1 condensates formed through overexpression.

**Figure 3.**
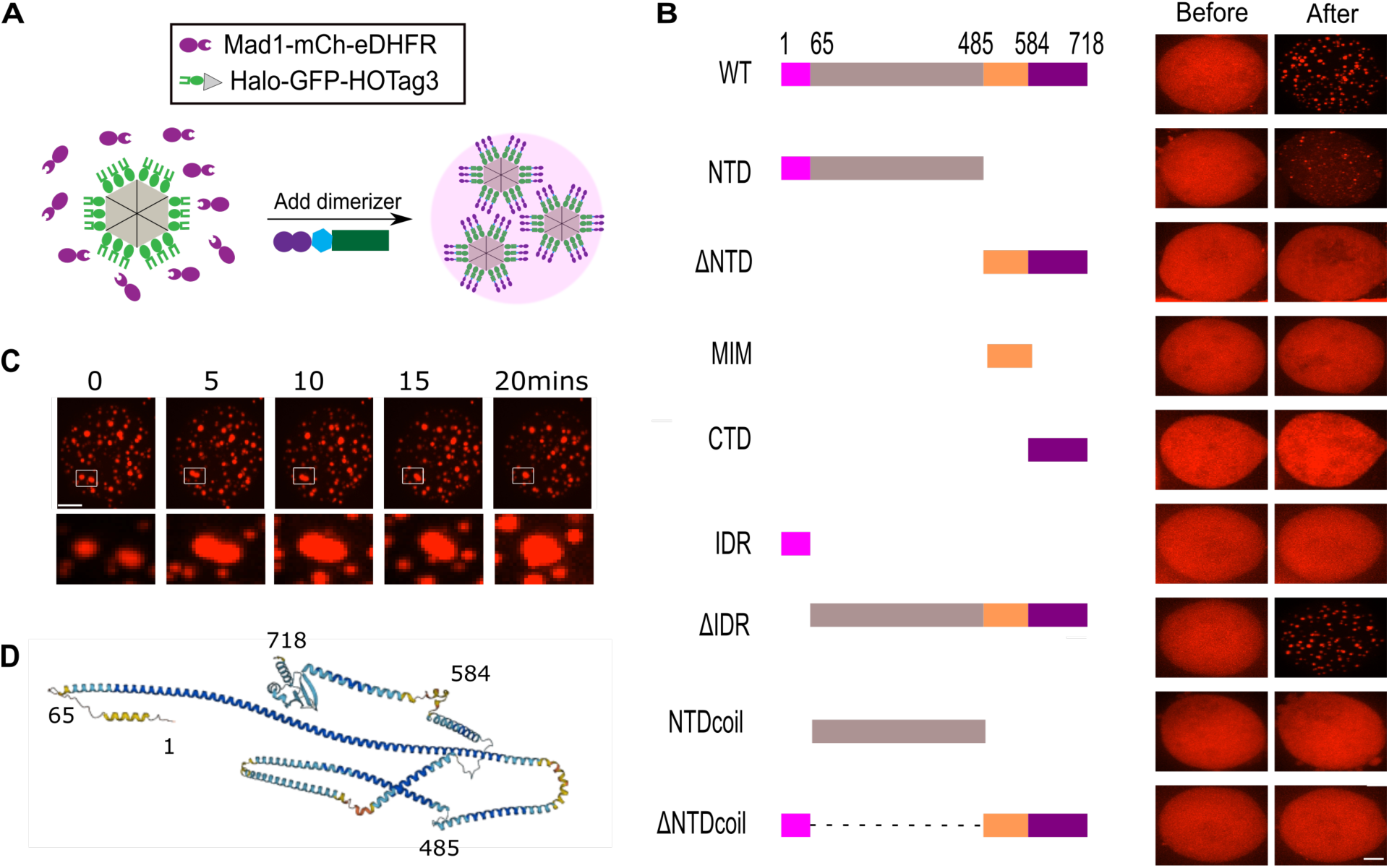
The coiled-coil domain within its N-terminus drives MAD1 phase separation. (A) Schematic to induce MAD1 phase separation with an oligomer seed using the chemical dimerization system. (B) Images for MAD1 truncations transfected in HeLa cells before and after adding dimerizers. (C) Fusion of MAD1 full-length condensates formed after adding the dimerizer. (D) MAD1 structure predicted by AlphaFold. Scale bar, 5 μm.

Functionally, MAD1 is divided into three domains (Fig. 3B): the N-terminus (NTD, 1-485), which aids in kinetochore localization; the MAD2 interaction motif (MIM, 485-584), which interacts with MAD2; and the C-terminus domain (CTD, 584-718), which binds to other checkpoint proteins (4). We observed that the NTD was sufficient to drive MAD1 phase separation, although with a reduced propensity (Fig. 3B). This suggests that the NTD is the primary driver for MAD1 phase separation, while the CTD and MIM also play a role. Further supporting the significance of the NTD in MAD1 phase separation, when the NTD was deleted from the full-length protein (Λ1IDR), it failed to form condensates (Fig. 3B). Conversely, neither MIM nor CTD alone formed condensates (Fig. 3B), reinforcing that the NTD is the main contributor to MAD1 phase separation.

We proceeded to test which domain within the NTD (1–484) was necessary for MAD1 phase separation. Although the structure of the NTD has not been solved, it is predicted to be largely coiled coil (4). The region from 1-65 is identified as an intrinsically disordered region (IDR), with a small helical structure predicted by Alphafold but with low confidence (Fig. 3D). Therefore, we divided the NTD into IDR (1-65) and NTDcoil (66-484) to examine their relative contributions to MAD1 phase separation (Fig. 3B). Neither IDR nor NTDcoil alone was sufficient for phase separation (Fig. 3B), indicating that both IDR and NTDcoil are necessary for MAD1 phase separation. Additionally, MAD1 with the NTD coil deleted (NTDcoil) did not form condensates, while the deletion of the IDR (Λ1IDR) did (Fig. 3B), suggesting that NTDcoil is more critical than the IDR. Collectively, these data demonstrate that the dominant driver for MAD1 phase separation resides within the NTD, particularly in the NTDcoil, while the IDR and other domains also contribute.

### MAD1 ectopic condensation promotes mitotic slippage

So far, our results indicate that overexpressed MAD1 can undergo ectopic condensation. Next, we investigated whether this ectopic condensation contributes to mitotic slippage. First, we used live imaging to monitor MAD1-mNeonGreen overexpressed cells after arresting them in mitosis with nocodazole (Fig. 4A). Cells with overexpressed MAD1 but without ectopic MAD1 condensates remained in the mitotic state, as shown by the compacted chromosomes visualized with miRFP670-H2B (Fig. 4B, C). In contrast, about 60% of cells with ectopic MAD1 condensates escaped mitosis within five hours of imaging, as indicated by the decompaction of chromosomes (Fig. 4B, C). These results suggest that the presence of MAD1 ectopic condensates correlates with a higher likelihood of experiencing mitotic slippage.

**Figure 4.**
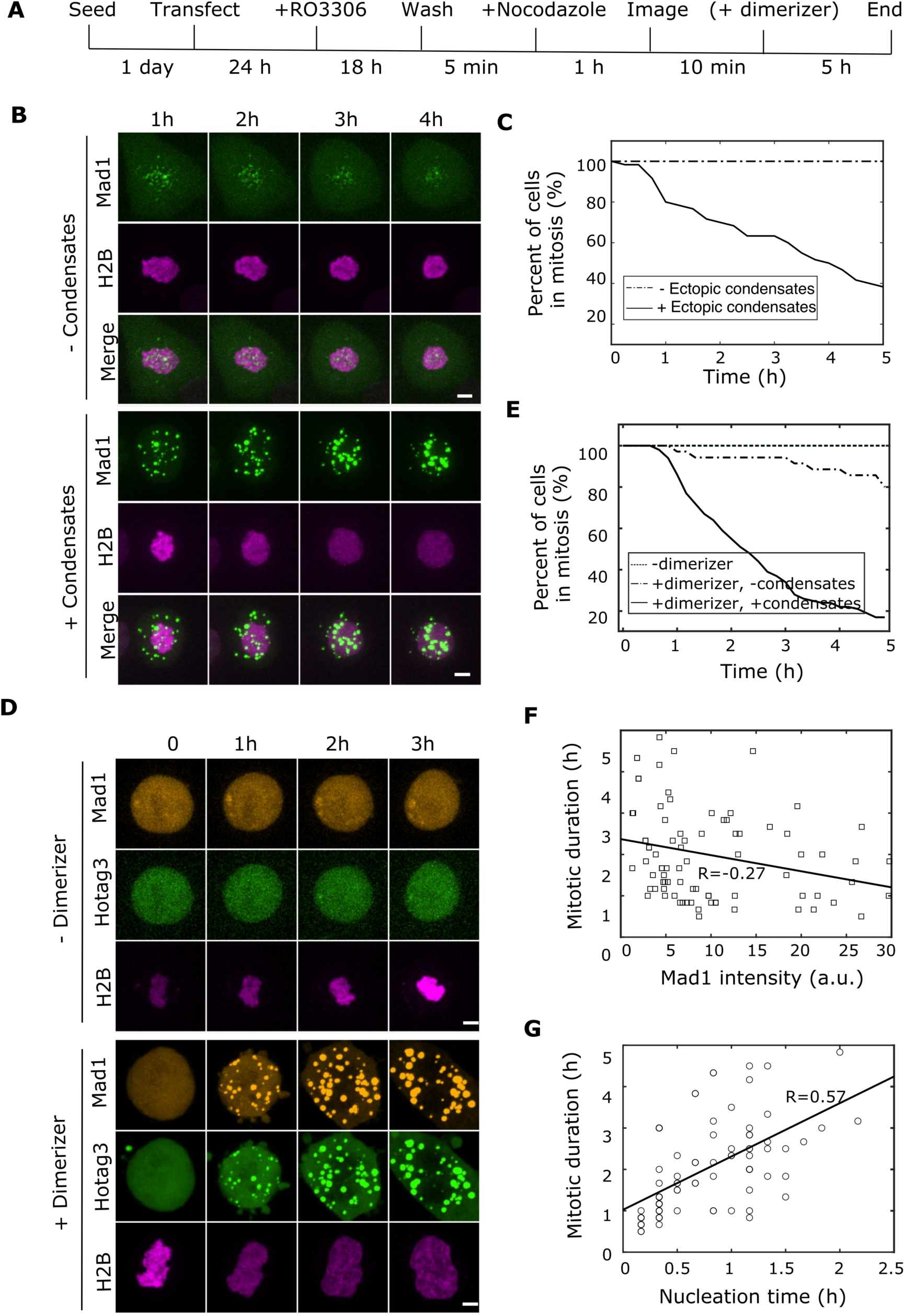
MAD1 ectopic condensation promotes mitotic slippage. (A) Schematic of the experimental procedure to follow mitotic slippage with live imaging. (B) Images and (C) quantification for the percentage of cells with/without ectopic MAD1 condensates that stayed in mitosis in HeLa cells transfected with MAD1-mNeonGreen and H2B-miRFP670. N= 57 cells from two replicates. (D) Images and (E) quantification for the percentage of cells that stayed in mitosis with/without adding dimerizers to induce MAD1 condensation by linking them to HOTag3. Halo-GFP-HOTag3 and H2B-miRFP670 were transiently transfected into HeLa cells stably expressing MAD1-eDHFR-mScarlet at low concentrations. N=132 cells from three replicates for + dimerizer and N=96 cells from three replicates for - dimerizer. (F) Mitotic duration with MAD1 intensity in the cells after adding the dimerizer. The solid line is a linear fit. Pearson correlation R = 0.27. (G) Mitotic duration and condensate nucleation time, the time when condensates appeared after adding the dimerizer, which represents the condensation propensity. The solid line is a linear fit. Pearson correlation R = 0.57. Scale bar, 5 μm.

However, cells with ectopic MAD1 condensates also have higher MAD1 concentrations. To decouple the contribution of MAD1 condensation from its concentration, we need to compare cells with similar MAD1 levels in both the condensed form and the diffusive form. To achieve this, we created a HeLa cell line expressing low levels of mCherry-eDHFR-MAD1 to prevent spontaneous ectopic condensation while still allowing induction of condensate formation through dimerization with the hexamer HOTag3 on demand (Fig. 3A). Similar to MAD1-mNeonGreen overexpression without ectopic MAD1 condensates (Fig. 4B, C), most MAD1-eDHFR-mScarlet cells maintained their mitotic states (Fig. 4D, E, Movie 2). Upon adding dimerizers, over 80% of these cells with MAD1 condensates exited mitosis (Fig. 4D, E, Movie 3), supporting the role of MAD1 ectopic condensation in mitotic slippage.

Although the MAD1-eDHFR-mScarlet stable cell line was used, minor differences in MAD1 expression at the single-cell level still existed. To determine whether the variation in MAD1 expression levels was responsible for the increased mitotic slippage, we plotted mitotic duration against MAD1 intensity in these cells and found little correlation (*R* = −0.27) (Fig. 4F). In contrast, a stronger correlation was observed (*R* = 0.57) when we plotted mitotic duration against the MAD1 condensate nucleation time (the time required for MAD1 condensates to form after the dimerizer is added, indicating the system’s propensity to phase separate) (Fig. 4G). These findings support the idea that MAD1’s ability to form ectopic condensates directly contributes to mitotic slippage.

### MAD1 ectopic condensates sequester MAD2 and Cyclin B1/CDK1

We investigated how MAD1 ectopic condensation affects mitotic slippage. Given that MAD1 ectopic condensates exhibit slow dynamics, we hypothesized that the sequestration of other mitotic proteins would impair their functions and result in a shorter mitotic duration. To test this, we first employed immunofluorescence (IF) to identify kinetochore proteins that co-localize with ectopic MAD1 condensates in mitotic cells. Using the CREST antibody, which recognizes various centromere proteins and is commonly used for detecting centromeres, we observed a co-localization signal with MAD1 ectopic condensates (Fig. 5A, C). This indicates that certain centromere proteins targeted by CREST are present in the ectopic MAD1 condensates.

**Figure 5.**
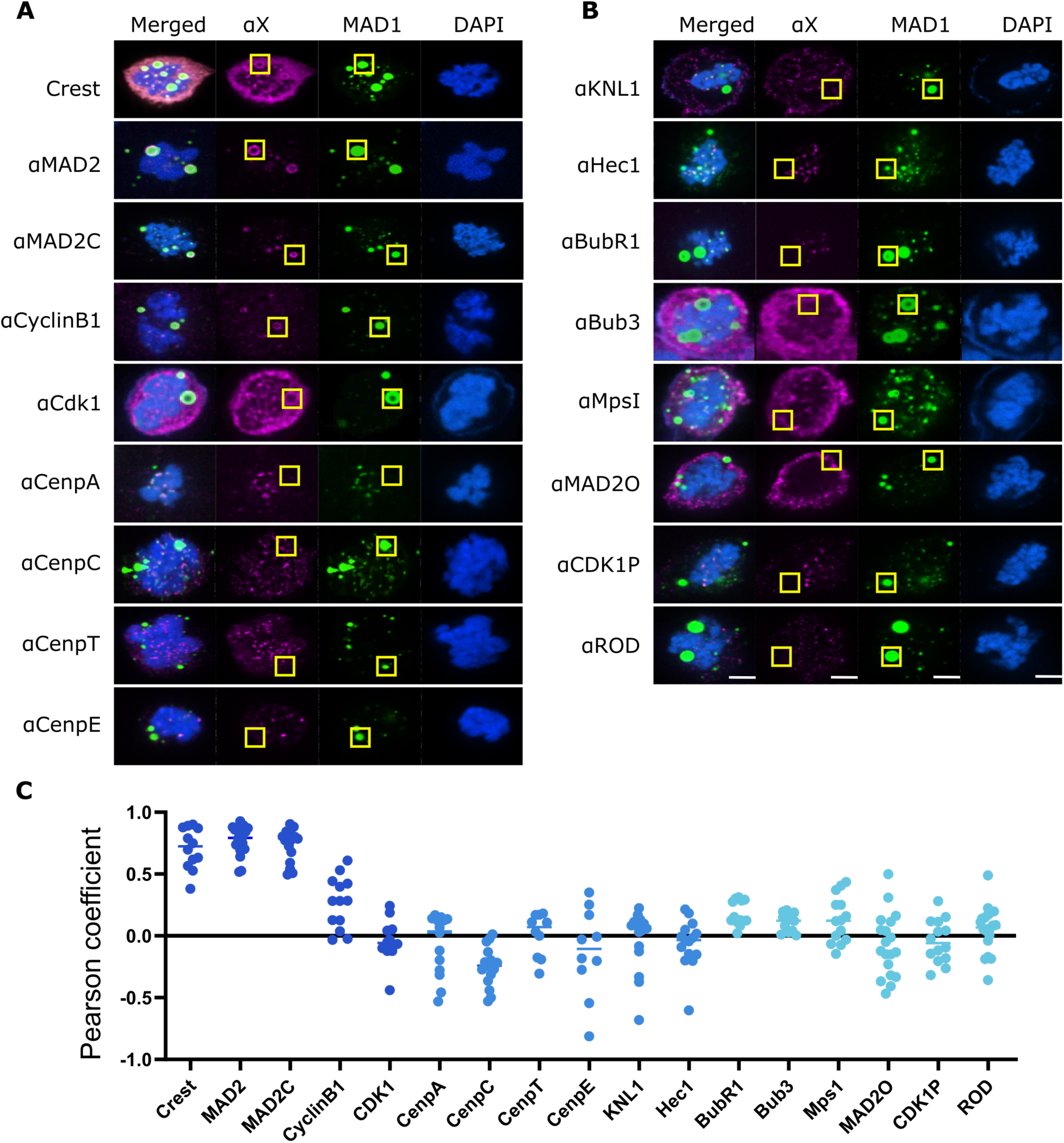
Co-localization of kinetochore proteins with MAD1 ectopic condensates. (A) Immunofluorescence of inner kinetochore and checkpoint proteins and their colonization with ectopic MAD1 foci. (B) Immunofluorescence of outer kinetochore proteins and their colonization with ectopic MAD1 foci. (C) Pearson coefficients between the Mad1 ectopic foci and kinetochore protein channels. Coefficients were calculated using the yellow boxed regions in B and C that highlight ectopic MAD1 foci, defined as MAD1 foci that are not co-localized with DNA. MAD1 ectopic foci are formed by transfecting HeLa cells with MAD1-mNeonGreen. Scale bar, 5 μm.

We then examined centromere components, including CENP-A, CENP-C, CENP-E, and CENP-T; none of which localized to ectopic MAD1 condensates in mitotic cells (Fig. 5A, C).

Similarly, outer kinetochore proteins, including BubR1, Bub3, Hec1, KNL1, Rod, and Mps1, did not localize to ectopic MAD1 condensates (Fig. 5B, C). We then tested Cyclin B1 and MAD2, two proteins reported to interact with MAD1 (4, 22). As expected, both Cyclin B1 and MAD2 localized to MAD1 ectopic condensates (Fig. 5A, C). Furthermore, CDK1, the master mitotic kinase that forms a complex with Cyclin B1 (23), was observed in ectopic MAD1 condensates (Fig. 5A, C), but not in its phosphorylated form (Fig. 5B, C). Using a MAD2 conformation-specific antibody, we observed Closed MAD2 in ectopic MAD1 condensates (Fig. 5A, C), but not Open MAD2 (Fig. 5B, C).

To confirm the IF results and rule out artifacts of fixed imaging, we expressed fluorescently labeled kinetochore proteins and imaged their localization with ectopic MAD1 condensates with live imaging (Fig. S2). Consistent with the IF results, Cyclin B1 and MAD2 localized to ectopic MAD1 condensates, whereas other proteins, including Mps1, CENP-A, CENP-C, CENP-T, KNL1, Spc24, Spc25, Nuf2, and Mis12, did not. Together, these findings suggest that MAD1 ectopic condensation might disrupt mitosis by affecting MAD2 and Cyclin B1/CDK1 functions.

### MAD1 condensation induces mitotic slippage via promoting Cyclin B1 degradation

We then examined whether the sequestration of Cyclin B1/CDK1 in the ectopic MAD1 condensates contributed to mitotic slippage. Cyclin B1, in complex with CDK1, is necessary to activate CDK1, allowing cells to remain in mitosis (24). One of the functions of the SAC is to prevent the degradation of Cyclin B1, ensuring that CDK1 remains active. However, inhibiting CDK1 activity with a small molecule while blocking Cyclin B1 degradation can lead to mitotic exit (25). This suggests that CDK1 activity, rather than the Cyclin B1 level itself, is essential for maintaining mitosis. Thus, the sequestration of Cyclin B1/CDK1 may promote mitotic slippage by either influencing Cyclin B1 degradation through the SAC or impacting CDK1 activity independent of Cyclin B1 levels.

To test this, we first monitored the Cyclin B1 level in cells overexpressing MAD1. We created a HeLa cell line with stable expression of Cyclin B1-mScarlet and transfected these cells with MAD1-mNeonGreen and H2B-miRFP670. Three-color microscopy imaging of these cells enabled us to observe MAD1 ectopic condensation, mitotic slippage, and Cyclin B1 levels simultaneously in nocodazole-arrested cells (Fig. 6A). Cyclin B1 remained at its level in cells without ectopic MAD1 foci but not in those with ectopic MAD1 foci (Fig. 6B, D; Fig.S3), indicating that MAD1 ectopic condensation weakens the checkpoint, promoting Cyclin B1 degradation. Furthermore, Cyclin B1 in MAD1 ectopic foci also degraded gradually (Fig. 6B, D; Movie 4), suggesting that sequestering Cyclin B1 does not protect it from the proteasome.

**Figure 6.**
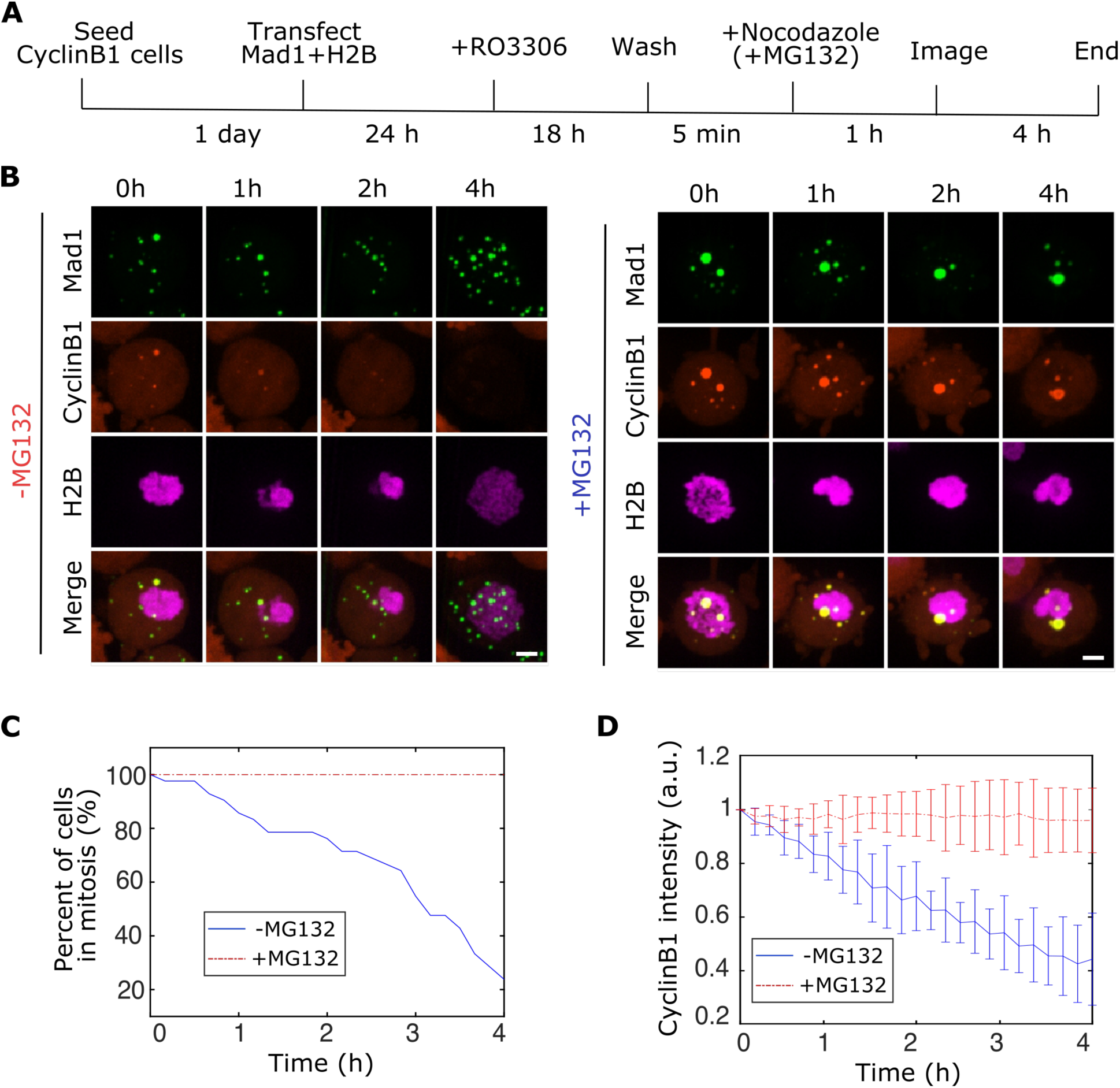
MG132 prevents MAD1 condensation-induced mitotic slippage. (A) The experimental procedure to visualize Cyclin B1 degradation and mitotic slippage in MAD1 overexpressed cells with or without the addition of MG132. Cyclin B1-mScarlet was stably integrated, while MAD1-mNeonGreen and H2B-miRFP670 were transfected into HeLa cells. (B) Representative images of mitotic Cyclin B1-mScarlet cells arrested with nocodazole that contain ectopic MAD1 condensates formed by overexpressed MAD1-mNeoGreen with/without MG132 added. H2B-miRFP670 was transfected to visualize chromosomes. (C) Quantification for percentage of cells exited mitosis for cells with ectopic MAD1 condensates. (D) Quantification for average Cyclin B1 intensity over time in cells with ectopic MAD1 condensates. Error bar, SEM. N=71 cells from two replicates for - MG132 and N=83 cells from two replicates for +MG132. Scale bar, 5 μm.

We then added the protease inhibitor MG132 to these cells to determine whether blocking Cyclin B1 degradation is sufficient to keep cells in mitosis. First, the addition of MG132 inhibited Cyclin B1 degradation (Fig. 6B, D, Fig. S3), demonstrating the effectiveness of MG132. In contrast to the results observed with the CDK1 inhibitor (25), MAD1 condensation-induced mitotic exit was prevented by MG132 (Fig. 6A, C, Fig. S3, Movie 5), suggesting that CDK1 activity is not inhibited by the sequestration of Cyclin B1/CDK1 into the MAD1 ectopic condensates. To rule out the effect of Cyclin B1 overexpression, we added MG132 to cells without Cyclin B1 overexpression and found that MAD1 foci-induced mitotic slippage was also prevented (Fig. S4). Therefore, we conclude that MAD1 condensation causes mitotic slippage not by sequestering Cyclin B1/CDK1 to directly inhibit CDK1 activity but mainly by weakening the checkpoint to promote Cyclin B1 degradation through Mad2 sequestration.

### MAD1 ectopic condensates trap MAD2 to weaken the checkpoint

Finally, we asked how MAD2 sequestration in the ectopic MAD1 condensates influenced the mitotic checkpoint. To this end, we expressed the Mad2 interaction motif (MIM) of MAD1 in cells and localized it, thus MAD2, to a synthetic condensate to investigate whether MAD2 sequestration would shorten the mitotic duration. This strategy was selected because mutating MIM in MAD1 could not separate the effects of MAD2 binding from MAD2 sequestration in the ectopic MAD1 condensates.

The synthetic condensates were made by fusing the disordered RGG-rich region of the Laf-1 protein (26) with Ferritin (27), a 24mer known for its efficiency in nucleating condensates for disordered proteins (Fig. 7A). We then utilized our chemical dimerization system to recruit the MIM to the RGG condensates. Since the high expression of MIM weakens the checkpoint by occupying the free open form of MAD2 (28), we made a HeLa cell line expressing a low level of MIM under doxycycline control. We verified that most cells sustained mitotic states in the presence of nocodazole during the four hours of imaging (Fig. 7B, C).

**Figure 7.**
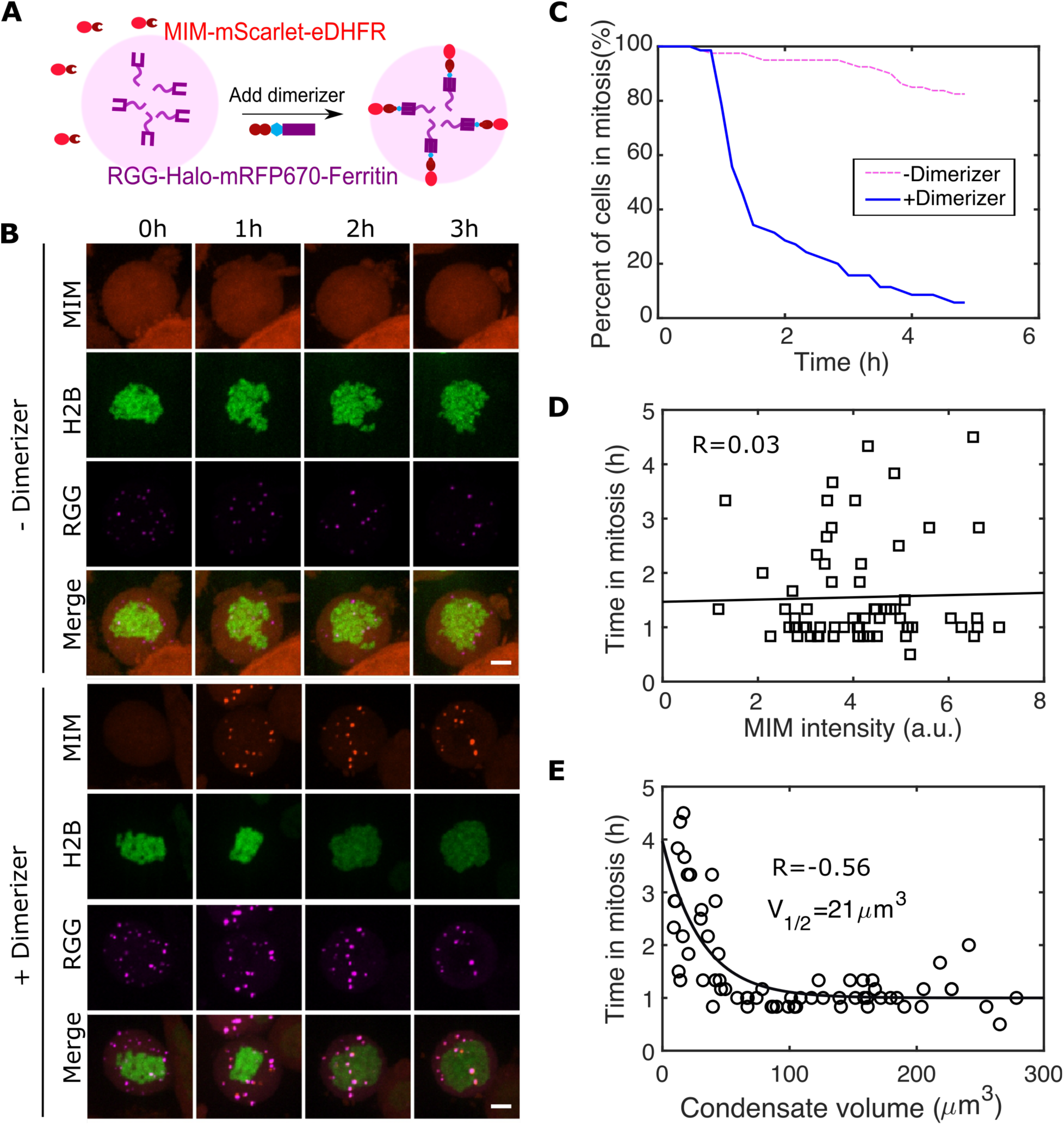
Sequestration of MAD2 to condensates contributes to premature mitotic exit. (A) Schematic to recruit MIM to synthetic condensates formed with RGG fused to oligomer Ferritin. (B) Representative images of MIM cells expressing RGG condensates with/without adding dimerizer after the first time point (C) Quantification of cells exited mitosis for cells with/without dimerizer. Dox inducible MIM-mScarlet-eDHFR was integrated, while RGG-Halo-miRFP670-Ferritin and H2B-GFP were transfected into HeLa cells. N=80 cells from two replicates for - dimerizer and N=82 cells from two replicates for + dimerizer. (D) Mitotic duration against MIM intensity for cells with MIM recruited to RGG condensates. The solid line is a linear fit. Pearson correlation R=0.03. (E) Mitotic duration against the volume of the RGG condensates for cells with MIM recruited to RGG condensates. The solid line is an exponential fit, yielding V_1/2_=21 μm^3^. Pearson correlation R = −0.56. Scale bar, 5 μm.

With the addition of the dimerizer, MIM was recruited to the RGG condensates (Fig. 7B). MAD2 was also observed in the RGG condensates (Fig. 8A), confirming the effectiveness of MIM in localizing MAD2 to these condensates. Adding the dimerizer significantly increased the fraction of cells exiting mitosis (Fig. 7B, D, Movie 6), suggesting that the sequestration of MAD2 to condensates contributes to mitotic slippage. In cells expressing RGG that did not form condensates, adding dimerizers did not induce slippage (Fig. S5, Movie 7), indicating that dimerizing MIM to RGG is not a significant cause of mitotic slippage. Moreover, for cells with MIM sequestered in RGG condensates, the mitotic duration showed little correlation (*R=0.03*) with MIM intensity in the cell (Fig. 7D). However, it negatively correlates (*R=-0.56*) with the volume of the RGG condensates (Fig. 7E). Interestingly, this relationship is not linear but rather exponential (Fig. 7E), suggesting that the effect of the condensation diminishes as the condensate volume increases, likely due to the limited availability of MAD2 to sequester.

**Figure 8.**
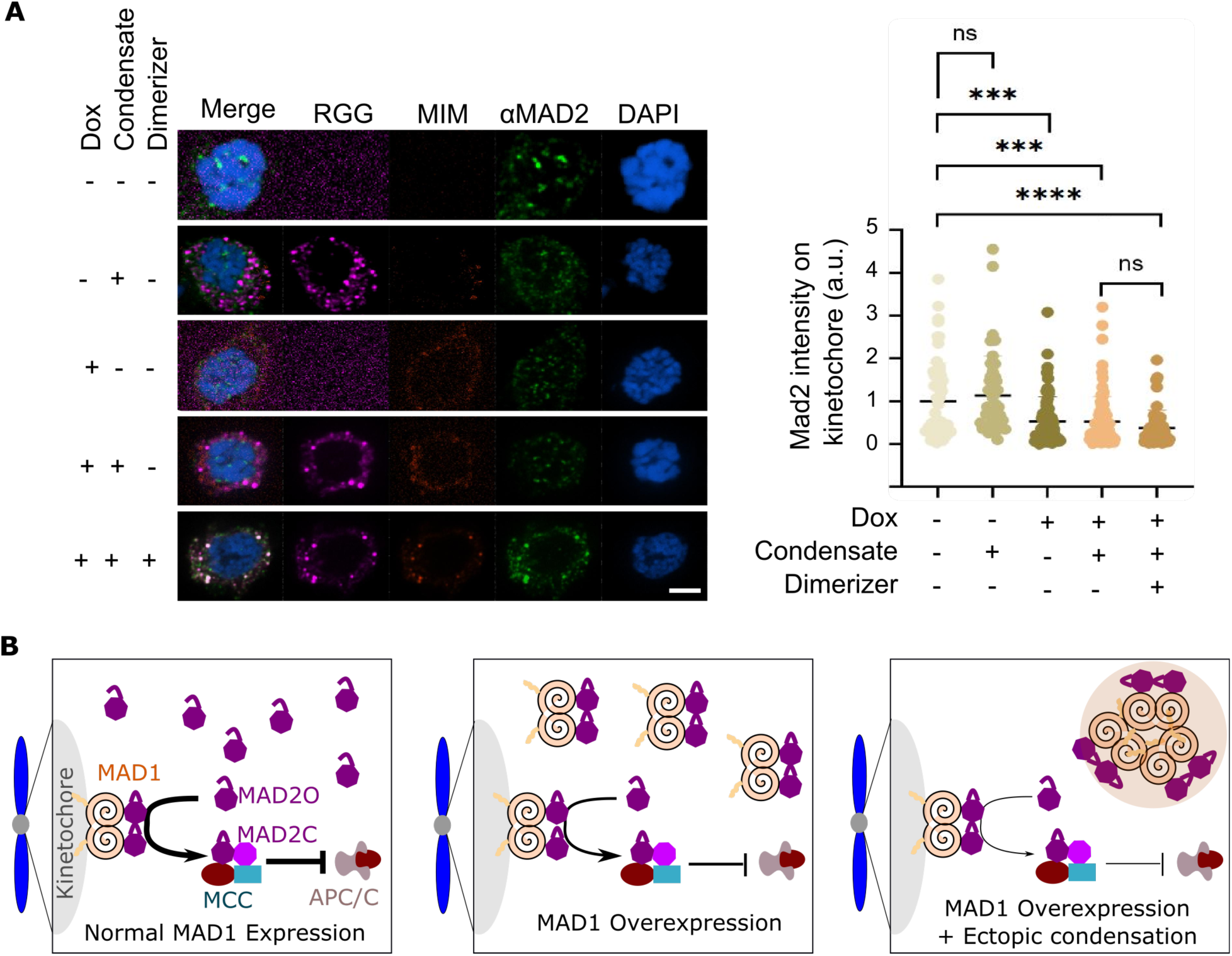
MAD1 ectopic condensation weakens the checkpoint without further reducing MAD2 localization on the kinetochore. (A) Representative images (left) and quantification (right) of MAD2 immunofluorescence on the kinetochore. Dox-inducible MIM-mScarlet-eDHFR was integrated into HeLa cells. RGG-Halo-miRFP670-Ferritin was transfected into the MIM cell line. MAD2 on kinetochores were identified as MAD2 foci that co-localized with the DAPI signal. Each dot represents one cell. Scale bar, 5 μm. *, p<0.01. ***, p<0.0001, ns, not significant. Two-tailed T test. (B) Working model. In normal cells (left), MAD2 is expressed more than MAD1. The MAD1-free MAD2 is in an open conformation (MAD2O). MAD2O is recruited to unattached kinetochores and converted to the closed form (MAD2C) to form the mitotic checkpoint complex (MCC) that inhibits the anaphase-promoting complex (APC/C) to prevent anaphase progression. When MAD1 is overexpressed (middle), MAD1 can occupy the free MAD2O, disrupting MAD2’s function to weaken the checkpoint. Overexpressed MAD1 undergoes ectopic condensation (right), further dampening the checkpoint by trapping MAD2 in the ectopic condensates.

Furthermore, MIM expression diminished MAD2 localization on the kinetochore in nocodazole-treated cells (Fig. 8A), which is consistent with the prior finding that MAD1 overexpression reduces MAD2 kinetochore localization (13). However, the addition of dimerizers did not further decrease MAD2 kinetochore localization (Fig. 8A). These results lead us to conclude that MAD1 ectopic condensates trap the diffusive form of MAD2, further weakening the checkpoint without worsening MAD2 kinetochore localization.

## Discussion

A robust mitotic checkpoint is essential for ensuring fidelity in chromosome segregation. It is also crucial for cancer therapies that rely on prolonged mitotic checkpoints to arrest cells in mitosis to induce mitotic cell death (29). However, mutations in checkpoint proteins often compromise the strength of the checkpoint, promoting cancer initiation and leading to cancer drug resistance via mitotic slippage (30). Previous studies have revealed that MAD1 overexpression weakens the checkpoint and promotes mitotic slippage (12, 13). In this work, we demonstrate that the formation of ectopic foci by overexpressed MAD1 also contributes to premature mitotic exit (Fig. 8B). We observe that the endogenous level of MAD1 has a low propensity to undergo phase separation, though whether MAD1 exists in a condensed form when concentrated at unattached kinetochores is unknown. When overexpressed, MAD1 can undergo global phase separation to form condensates in the cytosol of mitotic cells. These ectopic condensates sequester MAD2 without further reducing MAD2 localization on kinetochores beyond MAD1 overexpression, leading to rapid degradation of Cyclin B1 and triggering mitotic slippage. The trapping of MAD2 in the condensates likely limits the availability of diffusive MAD2 needed for efficient conversion from MAD2O to MAD2C. Therefore, targeting MAD1 condensation has the potential to reduce mitotic slippage in cancer cells that overexpress MAD1.

Condensate formation relies on multivalent interactions between biomolecules. While intrinsically disordered regions (IDRs) have been shown to promote phase separation for many proteins, increasing evidence suggests that coiled-coils are also significant drivers. Notable examples include SPD5 in the centrosome (30), the LINE-1 ORF1 protein (31), amphipathic helices enriched in proteins that condense to attenuate apoptosis (32), and engineered coiled coils (33). Coiled-coil domains have also been demonstrated to be sufficient to drive protein phase separation in molecular simulations (34). Our data indicate that the coiled-coil in MAD1 N-terminal domain (NTD) facilitates MAD1 condensation, adding another instance of coiled coil-driven phase separation. The mechanisms by which the coiled-coils contribute to MAD1 condensation remain to be investigated. MAD1 is predicted to exist as a dimer, with the dimerization domains localized at both the N-terminus and C-terminus (4). Our data suggest that both the small IDR domain and the coiled-coil at the N-terminus are essential for MAD1 phase separation. This implies that by promoting dimerization, the coiled-coil may enhance the IDR’s ability to drive condensation. It is also possible that the dimerized coils contribute to valence by interacting with charged residues on the surface. Alternatively, overexpressed MAD1 may fail to dimerize and thus interact with one another through hydrophobic residues. The way the IDR engages with the coiled-coil domain to promote MAD1 phase separation remains unknown but warrants investigation, as many phase separation proteins rely on cooperation between IDRs and coiled coils to fine-tune their phase behavior.

Phase separation is implicated in various cell functions, yet establishing a causal relationship has proven challenging. Traditional methods involve mutating the phase separation domain and/or substituting it with one from a different protein. However, these methods struggle to eliminate the possibility that other functions of the mutated domain aside from phase separation contribute to the observed effects. An alternative approach is to compare protein function with and without inducing phase separation. Notable examples include utilizing rapamycin-induced condensation to assess the enhancement of SUMOylation activity in condensates (35) and employing lenalidomide-induced Myc phase separation to evaluate its effect on transcription (36). In this study, we applied a similar method to establish a causal relationship between MAD1 condensation and mitotic slippage.

Ectopic condensates are present in many cancers, so understanding how these condensates contribute to cancer initiation and progression holds promise for cancer prevention and therapy (37–39). Because biomolecular condensates are linked to cellular functions through various mechanisms (40, 41), the roles of ectopic condensates in cancer biology can also be diverse. In leukemias, for instance, the fusion protein NUP98-HOXA9 undergoes phase separation due to the intrinsically disordered domain (IDR) in NUP98; this phase separation contributes to oncogenesis by forming aberrant chromatin looping that upregulates the expression of the target genes of the transcription factor HOXA9 (42). In cancer cells that use the alternative lengthening of telomeres (ALT) pathway for telomere maintenance, the coalescence of mis-localized PML bodies on ALT telomeres pulls telomeres together to provide repair templates for homology-directed telomere DNA synthesis in ALT cancer cells (43). Our work illustrates how sequestration by ectopic condensates results in the loss of protein function, contributing to an array of mechanisms that aberrant phase separation in cancer cells employ to disrupt cellular functions.

## Materials and Methods

### Plasmids

The MAD1-mNeogreen plasmid was gifted by Dr. Beth Weaver (13). Plasmids containing eDHFR and Halo for cell imaging are derived from our previous plasmids containing a CAG promoter obtained from E. V. Makeyev (43, 44). RGG is from a plasmid gifted by Benjamin Schuster(45). Ferritin is from Addgene (27). Cyclin B1-GFP is from Addgene (26061), GFP-Mps1 is from Addgene (63702), CENP-A, CENP-C, CENP-T, KNL1, Spc24, Spc25, Nuf2, Mis12 were obtained from Dr. Michael Lampson, HOTag3 was ordered as complementary oligomers (IDT), annealed, and cloned using T4 ligation (NEB M0202S). The plasmids are transfected into cells using Lipofectamine 2000 or Lipofectamine 3000 (Invitrogen) following manufacturer protocol.

### Cell line and cell culture

Unless otherwise noted, experiments were performed with HeLa cells obtained from E. V. Makeyev (44). The HeLa MAD1-mNeon CRISPR knock-in cell line was obtained from Ajit P. Joglekar (16). The LacO cell line was obtained from Dr. Michael Lampson. The U2OS cell line with PML deleted were gifts from Dr. Eros Lazzerini Denchi (46). HeLa cells with endogenous PML tagged with Clover (U2OS Clover-PML) were obtained from Graham Dellaire (47). Stable cell lines were created with the RMCE system, as reported previously(48, 49). Briefly, HeLa acceptor cells at 60–80% confluency in a well of a six-well plate were transfected with 1 μg of plasmid with the targeted gene and 10 ng of a Cre recombinase plasmid using Lipofectamine 2000. Two days after transfection, 1 μg/mL puromycin was added to the growth medium to select stable cell lines.

Cells were cultured in a growth medium (Dulbecco’s Modified Eagle’s medium with 10% FBS and 1% penicillin–streptomycin) at 37 ^°^C in a humidified atmosphere with 5% CO_2_. For mitotic cell arrest, cells were first treated with 9 μM CDK1 inhibitor (RO-3306, cat# SML0569, Sigma-Aldrich) for 16 hours, washed for 5 mins, and then treated with 300 ng/ml of nocodazole.

### Protein dimerization with chemical dimerizers

The dimerizer TFH (**T**MP-**F**luorobenzamide-**H**alo) was used. Its synthesis and storage were previously reported (19). For live imaging, the dimerizers were first diluted to 200 nM in growth medium and then added to the imaging chamber to a final working concentration of 100 nM after the first round of imaging.

For immunofluorescence (IF), TFH was added directly to the growth medium to a final working concentration of 100 nM after 30 minutes of adding nocodazole. Cells were incubated for 1 hour before fixation.

### Cell imaging and image processing

For live imaging, cells were seeded on 22 x 22 mm glass coverslips coated with poly-D-lysine (Sigma-Aldrich). When ready for imaging, coverslips were mounted in magnetic chambers (Chamlide CM-S22-1, LCI) with cells maintained in growth medium at 37 °C in an environmental chamber (TOKAI HIT Co., Ltd.). Images were acquired with a microscope (ECLIPSE Ti2) with a 60 × 1.4 NA objective, a 16 XY Piezo-Z stage (Nikon Instruments Inc.), a spinning disk (Yokogawa), an electron multiplier charge-coupled device camera (IXON-L-897), and a laser merge module that was equipped with 488 nm, 561 nm, 594 nm, and 630 nm lasers controlled by NIS-Elements Advanced Research (Nikon Instruments Inc.). For live imaging, images were taken with 1 μm spacing between Z slices, for a total of 20 μm. For movies, images were taken at 10-minute intervals for up to 5 hours. For fixed imaging, images were taken with 0.3 μm spacing between Z slices, for a total of 8 μm.

For FRAP, MAD1 foci were bleached with a 405 nm laser at 40% power for 40 ms on the Nikon confocal microscope, and images were taken every 10 seconds before and after bleaching.

Images were processed and analyzed using NIS Elements Software (Nikon Instruments Inc.). Maximum projections were created from z stacks, and thresholds were applied to the resulting 2D images to segment and identify MAD1 and synthetic condensate foci. The quantifications were exported to Excel and plotted in MATLAB (Mathworks).

### MAD1 saturation concentration calculation

Mitotic cells were manually annotated as ROIs to quantify overexpressed MAD1-mNeon in HeLa cells. After background subtraction, the average mNeon fluorescent intensity in the ROI was obtained (I_Over). To convert the fluorescent intensity to protein concentration, we calculated the mean mNeon fluorescent intensity in the MAD1-mNeon CRISPR knock-in mitotic cells (I_Crispr). By taking the reported endogenous MAD1 concentration in Hela cells as 20 nm and the CRISPR labeling efficiency as 72% (16, 17), we estimated the concentration of overexpressed Mad1 (C_Over) following C_Over=I_Over/I_Crispr*72%*20 nm.

To calculate saturation concentration, cells were visually classified as having either ectopic or no ectopic MAD1 foci. MAD1 concentration in cells was plotted against the condensate formation and fitted with a logistic regression line plus 95% confidence intervals using Prism (GraphPad). The midpoint concentration between foci- and non-foci-containing cells was reported as the saturation concentration along with standard error.

### Immunofluorescence

Cells were grown on 12 mm circle cover glass (Fisher 12541001), washed once with room temperature PBS, then permeabilized in 37°C PHEM-T buffer (60mM PIPES, 25mM HEPES, 10mM EGTA, 1mM MgCl_2_, 0.1% Triton X-100). Next, cells were fixed for 10 minutes at room temperature in PHEM-T buffer with 4% paraformaldehyde and then washed 3 times with room temperature PBS for 5 minutes per wash. Cells were blocked for 1 hour in a blocking buffer (1X PBS, 5% serum, 0.3% Triton X-100) at room temperature in a humidified box protected from light. Primary antibodies were diluted in blocking buffer and were applied to cover glass and stained overnight in a humidified box protected from light at 4°C. Cells were next washed 3 times with room temperature PBS for 5 minutes per wash, secondary antibodies (Alexa fluor 488 or 647) were diluted (1:1000) in antibody dilution buffer(1X PBS, 1% BSA, 0.3% Triton X-100), and applied to cells for 1 hour in a humidified box protected from light at room temperature. Cells were washed 3 times with room temperature PBS for 5 minutes per wash, stained with DAPI (1 ng/mL) for 1 minute, and then mounted onto glass slides with mounting media (SouthernBiotech 0100-01).

Primary antibodies used and their dilutions are: CREST(Immunovision HCT-0100) 1:5000, MAD2((107-276-3), Santa Cruz Sc-65492) 1:300, Cyclin B1((GNS1), Santa Cruz Sc-245) 1:500, CENP-A((3-19), ThermoFisher MA120832) 1:1000, CENP-C(ThermoFisher PA566154) 1:1000, CENP-E(ThermoFisher MA518025) 1:500, CENP-T (ThermoFisher PA583540) 1:1000, KNTC1((ROD), ThermoFisher PA555234) 1:1000, Bub3(BDBiosciences 611730) 1:100, Cdc2((POH1), Cell Signaling 9116) 1:300, Phospho-cdc2((Tyr15)(10A11), Cell Signaling 4539) 1:100, KNL1(generous gift from Iain Cheeseman) 1:1000. Following antibodies generously gifted from Jennifer G. DeLuca and Jakob Nilsson: BubR1(0.85 µg/mL), MAD2-C (1.6 µg/mL), Hec1(1.0 µmL), MAD2-O (4.0 µg/mL).

### Statistical methods

All error bars represent means ± SEM. Statistical analyses were performed in MATLAB (Mathworks) and Prism (GraphPad). Detailed statistical methods were described in figure legends. Statistical significance: N.S., not significant, P > 0.05; *P < 0.05; **P < 0.01; ***P < 0.001.

## Acknowledgments

We thank Beth Weaver, Ajit Joglekar, Iain Cheeseman, Jennifer G. DeLuca, Jakob Nilsson, Eros Lazzerini Denchi, and Graham Dellaire for gifting reagents. HZ acknowledges support from the National Science Foundation (NSF MCB-2145083), JJ acknowledges support from the National Institutes of Health T32 Fellowship (T32GM133353), and DC acknowledges support from the National Institutes of Health (GM118510).

## Author Contributions

H.Z. conceptualized the study. J.T., J.J., and H.Z. performed experiments, analyzed data, and made figures. R.L. and D.C. prepared the dimerizers. H.Z. wrote the manuscript with comments from other authors.

## Data and Materials Availability

All data needed to evaluate the conclusions in the paper are present in the paper and the Supplementary Materials.

## Supplemental information For

### Movie 1-7

**Movie 1. Ectopic MAD1 foci in mitotic cells.** HeLa cells expressing MAD1-mNeoGreen and miRFP670-H2B were arrested in mitosis with nocodazole. Scale bar, 5 μm.

**Movie 2. Overexpressed MAD1 in diffusive form does not cause mitotic slippage.** HeLa cells expressing mCherry-eDHFR-MAD1, Halo-GFP-HoTag3, and miRFP670-H2B were arrested in mitosis with nocodazole. No dimerizer was added. Scale bar, 5 μm.

**Movie 3. Overexpressed MAD1 in condensed form causes mitotic slippage.** HeLa cells expressing mCherry-eDHFR-MAD1, Halo-GFP-HOTag3, and miRFP670-H2B arrested in mitosis with nocodazole, dimerizer was added after the first timepoint. Scale bar, 5 μm.

**Movie 4. CyclinB degradation in cells with MAD1 ectopic condensates.** Cyclin B1-mScarlet stable HeLa cell line expressing MAD1-NeoGreen and miRFP670-H2B arrested in mitosis with nocodazole, without adding MG132. Scale bar, 5 μm.

**Movie 5. Prevention of CyclinB degradation in cells with MAD1 ectopic condensates using MG132.** Cyclin B1-mScarlet stable HeLa cell line expressing MAD1-Neogreen and miRFP670-H2B arrested in mitosis with nocodazole, MG132 was added together with nocodazole right after CDK1 inhibitor washout. Scale bar, 5 μm.

**Movie 6. Recruiting MIM to condensed RGG-Ferritin leads to mitotic slippage.** MIM-mScarlet-eDHFR stable HeLa cell line expressing RGG-Halo-miRFP670nano3-Ferritin condensates and GFP-H2B arrested in mitosis with nocodazole, dimerizer was added after first time point. Scale bar, 5 μm.

**Movie 7. Recruiting MIM to diffusive RGG-Ferritin does not lead to mitotic slippage.** MIM-mScarlet-eDHFR stable HeLa cell line expressing RGG-Halo-miRFP670nano3-Ferritin in diffusive form and GFP-H2B arrested in mitosis with nocodazole, dimerizer was added after first time point. Scale bar, 5 μm.

### Supplemental Figure 1-5

**Figure S1.**
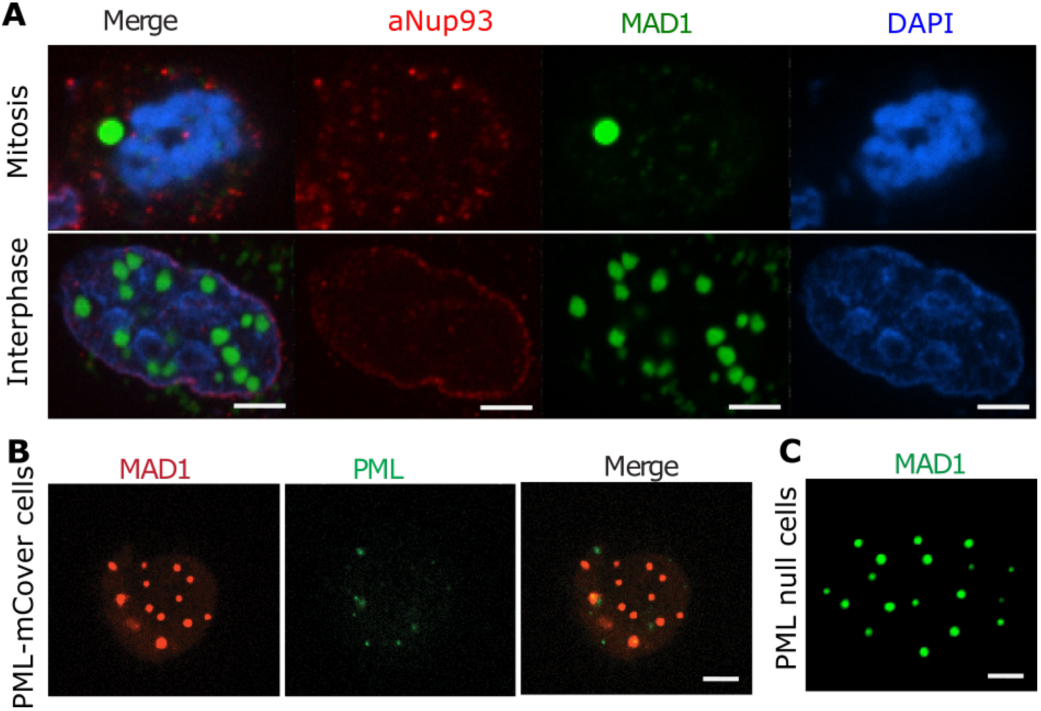
MAD1 ectopic foci do not co-localize with PML and Nup93 in mitotic cells. (A) Representative immunofluorescence images of Nup93 with MAD1 foci formed with overexpressed MAD1 in mitotic and interphase HeLa cells. (B) Representative live cell images of MAD1 foci formed with overexpressed MAD1 in mitotic U2OS cells with endogenous PML tagged with mClover via CRISPR. (C) Representative live cell images of MAD1 foci formed with overexpressed MAD1 in mitotic U2OS cells with PML deleted with CRISPR. Scale bar, 5 μm.

**Figure S2.**
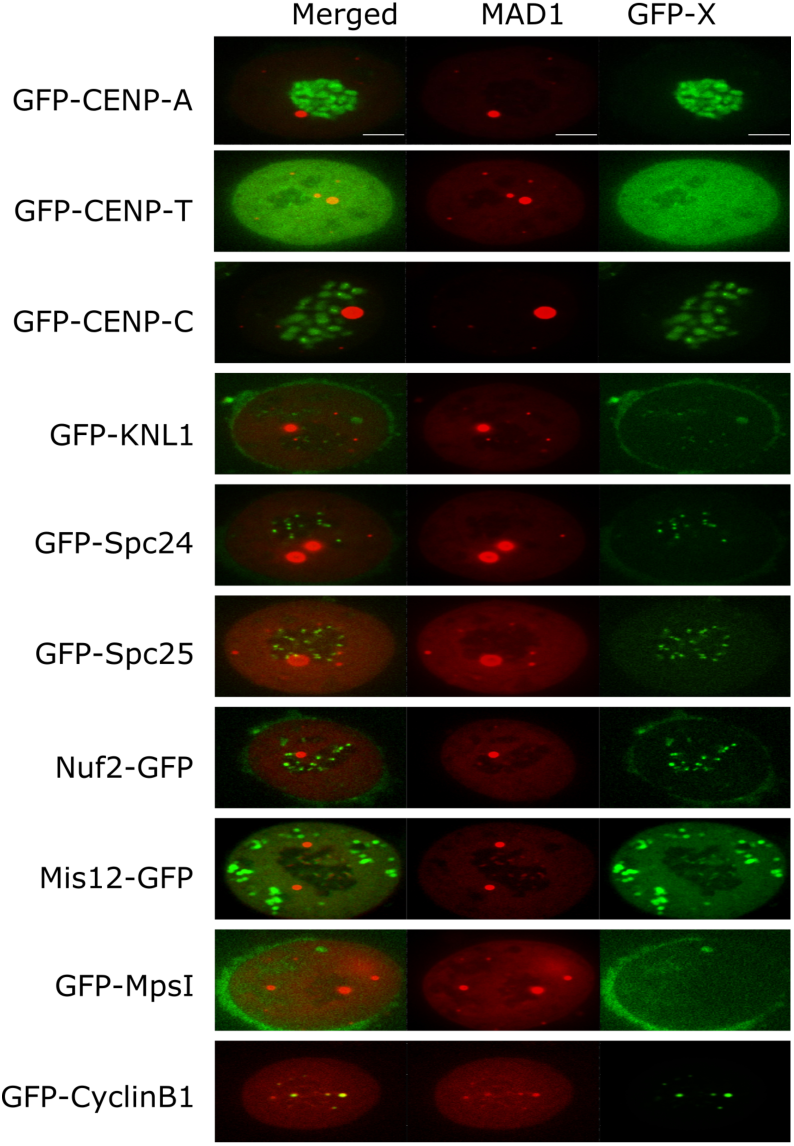
Live imaging of GFP-fused kinetochore proteins with ectopic MAD1 foci (red). HeLa cells were transiently transfected with indicated proteins. Scale bar, 5 μm.

**Figure S3.**
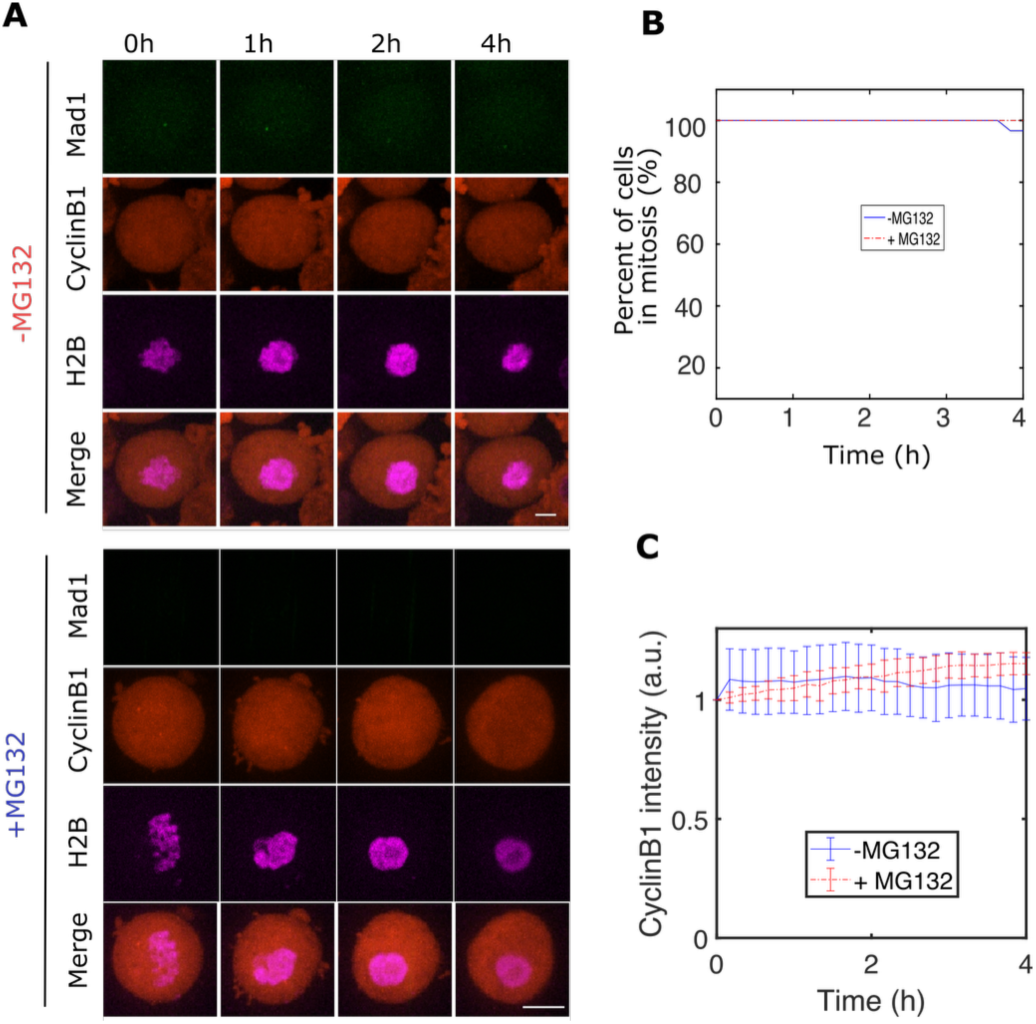
Cyclin B1 degradation in cells without MAD1 overexpression. (A) Representative images and (B) quantification for the percentage of cells that exited mitosis and (C) quantification for average Cyclin B1 intensity over time in mitotic cells. Hela cells with Cyclin B1-mScarlet stably expressed, MAD1-NeonGreen overexpressed but no ectopic condensates. Cells were arrested with nocodazole, with/without MG132 added. H2B-miRFP670 was transfected to visualize chromosomes. Scale bar, 5 μm.

**Figure S4.**
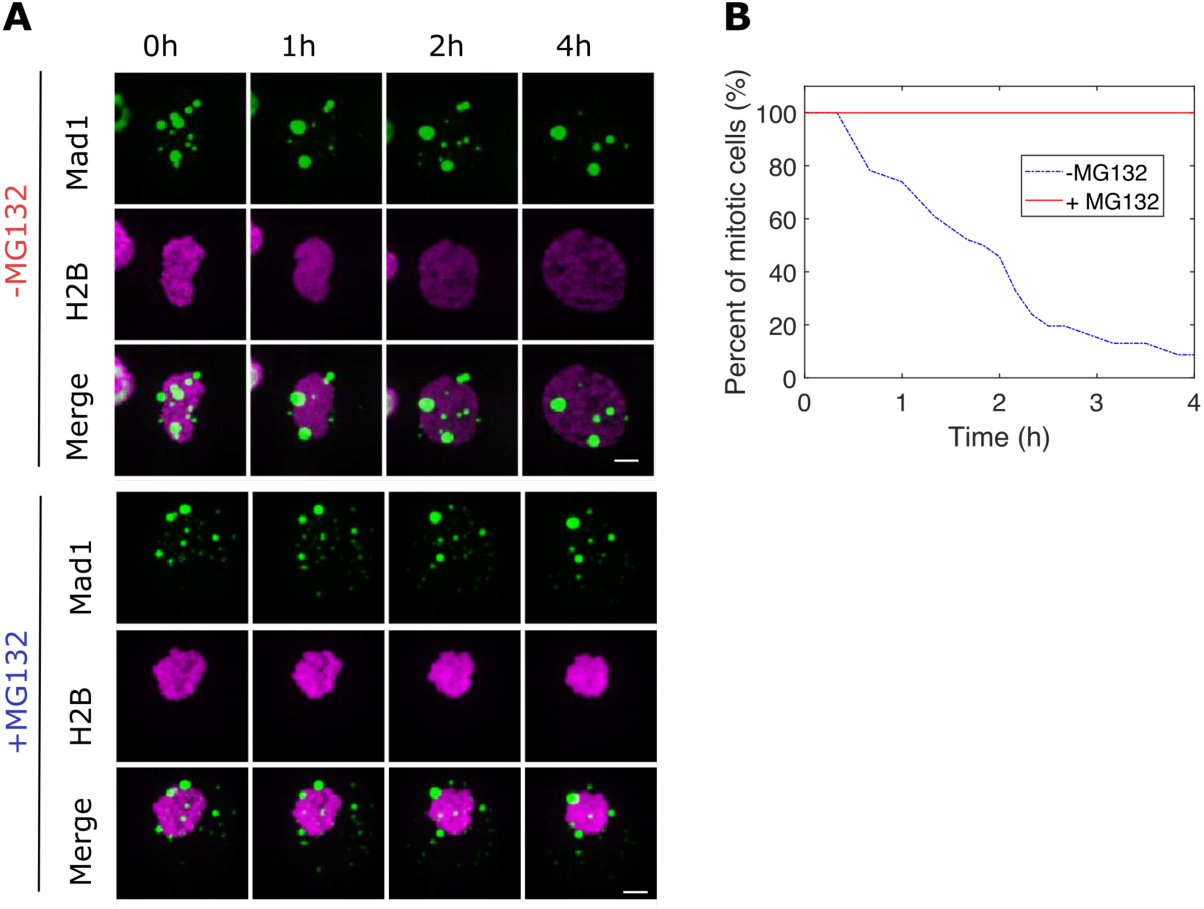
MG132 prevents mitotic slippage caused by MAD1 overexpression. (A) Representative images and (B) quantification for percentage of cells exited mitosis. HeLa cells with MAD1-mNeonGreen overexpressed and ectopic condensates formed but without Cyclin B1 overexpressed. Scale bar, 5 μm.

**Figure S5.**
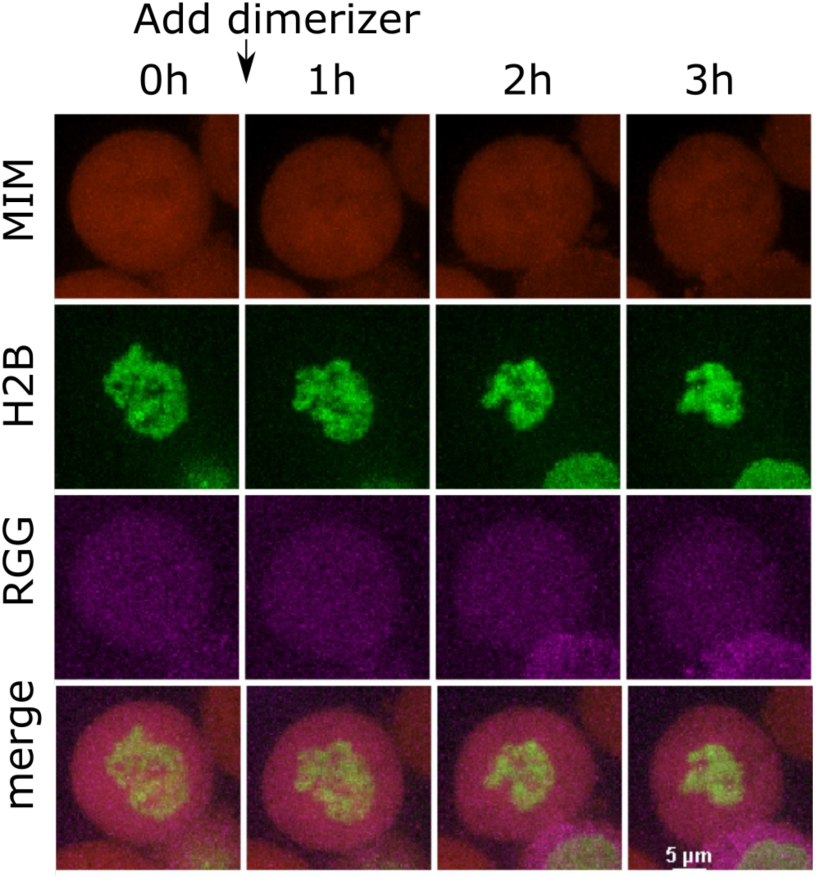
Dimerizing MIM to RGG-Ferritin in non-condensate form did not cause mitotic slippage. Representative images of MIM-mScarlet-eDHFR stable HeLa cell line expressing low levels of RGG-Halo-miRFP670-Ferritin that did not form condensates. The dimerizer was added at the second time point. Scale bar, 5 μm.

